# A Stable Reporter Cell Line to Study Respiratory Syncytial Virus NS2-Mediated Inhibition of *IFNB* Promoter Activation

**DOI:** 10.64898/2026.02.24.706909

**Authors:** Marta Acchioni, Lydie Martin, Elina Gerber-Tichet, Chiara Acchioni, Dacquin M. Kasumba, Marco Sgarbanti, Nathalie Grandvaux

**Affiliations:** Department of Infectious Diseases, Istituto Superiore di Sanità, viale Regina Elena 299, 00161, Rome, Italy; Centre de recherche du Centre Hospitalier de l’Université de Montréal (CRCHUM), Montréal, Québec, H2X0A9, Canada; Department of microbiology, infectiology and immunology, Faculty of medicine, Université de Montréal, Montréal, H3C 3J7, Canada; Department of biochemistry and molecular medicine, Faculty of medicine, Université de Montréal, Montréal, H3C 3J7, Canada

**Keywords:** Respiratory Syncytial Virus, NS2, Interferon, reporter system, reporter cell lines, luciferase, RIG-I

## Abstract

Human respiratory syncytial virus (RSV) is an enveloped, non-segmented, negative-sense RNA virus and a major cause of severe lower respiratory tract disease in infants, older adults, and immunocompromised individuals. A hallmark of RSV pathogenesis is early evasion of innate immunity mediated by the non-structural proteins NS1 and NS2. NS2 is a multifunctional interferon (IFN) antagonist that dampens both IFNβ induction and type I IFN responsiveness by targeting multiple nodes of the RIG-I/IFN signaling axis. Because NS2 inhibition is typically partial and context dependent, its quantitative assessment in conventional cell-based assays can be challenging. Here, we report the generation and characterization of a stable, virus-free reporter system in human lung epithelial A549 cells to measure NS2-mediated inhibition of RIG-I–dependent activation of the human IFNB promoter. The platform combines stable NS2 expression with genomic integration of a dual cassette encoding (i) a non-targeting short hairpin RNA that constitutively activates RIG-I signaling and (ii) a firefly luciferase reporter driven by the IFNB promoter. This configuration provides a sensitive and reproducible luminescence readout of IFNB promoter activity under controlled, infection-free conditions, enabling robust quantification of NS2 antagonism with reduced experimental variability. The luminescence-based format is readily scalable for early-stage discovery and prioritization of small molecules or biologics that counteract NS2 function. More broadly, the modular design can be adapted to other viral immune antagonists, signaling pathways, or cell types, supporting comparative studies where pathway modulation must be evaluated under highly standardized and reproducible conditions.

## Introduction

Human Respiratory Syncytial Virus (RSV), a respiratory enveloped, non-segmented negative sense RNA virus, remains a leading cause of severe lower respiratory tract infections in infants, the elderly, and immunocompromised individuals worldwide (1). The host’s first line of defense against viral infection relies on a rapid and coordinated type I interferon (IFN) response. Detection of viral dsRNA intermediates by pathogen recognition receptors (PRR) activates downstream signaling through interaction with the mitochondrial MAVS adaptor to activate downstream IκB kinase (IKK)-related family members, Tank Binding Kinase 1 (TBK1)/IKKε, ultimately leading to the phosphorylation and nuclear translocation of Interferon Regulatory Factor 3 (IRF3) and nuclear factor-kappa B (NF-κB) to transactivate the *IFNB* gene. Secreted IFNβ then binds to the type I IFN receptor (IFNAR1/IFNAR2), triggering the Janus kinase/signal transducers and activators of transcription (JAK/STAT) signaling cascade and the formation of the IFN-stimulated gene factor 3 (ISGF3) complex (STAT1, STAT2, and IRF9). ISGF3 translocates to the nucleus and binds IFN-stimulated response elements (ISREs) in the promoter of hundreds of IFN-stimulated genes (ISGs), establishing an antiviral state that limits viral replication and spread (2–4).

A key feature of RSV pathogenesis is its ability to evade this antiviral program through the activity of its non-structural (NS) proteins, NS1 and NS2, which are expressed early during infection (5, 6). The NS2 protein targets multiple nodes in the IFN pathway to inhibit IFNβ expression and responsiveness. Notably, NS2 suppresses *IFNB* transcription through binding to inactive RIG-I and MDA5, blocking the ubiquitination required for their activation; NS2 binding to RIG-I N-terminal Caspase activation and recruitment domain (CARD) disrupts RIG-I interaction with the MAVS adaptor, thereby preventing downstream signal transduction. Additionally, NS2 contributes to the degradation of STAT2, thereby blocking ISGs expression in response to IFNβ signaling (7–11). Through this dual inhibition of both IFNβ production and downstream responsiveness, NS2 significantly weakens the host antiviral defense and contributes to viral spreading and pathogenesis.

In recent years, major progress has been achieved in the prevention of RSV infections. New vaccines have been approved since 2023 including formulations for use in older adults and a vaccine specifically available for pregnant women, the latter aiming to protect newborns through maternal immunization (12, 13). However, vaccines for babies remain challenging with clinical trials halted by FDA in 2024 due to safety signals. Additionally, long-acting monoclonal antibodies such as Nirsevimab and Clesrovimab have shown strong efficacy in preventing severe RSV disease in infants and are now included in public health recommendations (14, 15). Although these preventive approaches represent major achievements in reducing RSV-associated disease burden, their impact may be constrained by rising vaccine hesitancy, uneven access across regions and populations, and attenuated immune responses in those at highest risk (infants, older adults, and immunocompromised individuals). Consequently, there remains a critical unmet need for therapeutic antiviral compounds capable of treating established RSV infection and suppressing viral replication and associated pathogenesis. Emerging research has identified NS2 as an attractive therapeutic target due to its central role in inhibiting host IFN responses (10, 16). Future development of small molecule inhibitors targeting NS2 would complement existing prophylactic strategies and address the critical therapeutic gap in RSV treatment.

In this study, we developed a reporter cell line using human lung epithelial A549 cells stably expressing the RSV NS2 protein, a luciferase reporter driven by the human *IFNB* promoter, and constitutively expressed RIG-I-activating dsRNA species. This virus-free cell-based system is suitable for quantifying NS2-mediated antagonism of *IFNB* induction in a controlled, reproducible manner suitable for quantifying RSV NS2-mediated inhibition of the IFNβ response.

## Materials and Methods

### Plasmids

The plasmid hNS2-pcDNA5, encoding a codon-optimized sequence of the RSV Long strain NS2, was previously described (8). To generate the hNS2-3xFlag-pCMV-3Tag1a construct, the hNS2 sequence was subcloned into the 3xFlag-pCMV-3Tag1a backbone (a kind gift from Dr. J. Archambault, McGill University). The pSIREN RetroQ IFN-inducer *IFNB* promoter luciferase retroviral vector was obtained by subcloning the IFN-inducer *IFNB* promoter luciferase cassette into the pSIREN RetroQ retroviral vector (Clontech Laboratories, Inc., now Takara Bio USA, Inc.) using the BamHI and EcoRI restriction sites. The IFN-inducer *IFNB* promoter luciferase cassette was generated by *de novo* gene synthesis (GenScript Biotech Netherlands BV) and is composed as follows: an IFN-inducer sequence previously described in (17), that encodes for a short hairpin RNA (shRNA) under the control of the pU6 RNAPol III promoter present in the pSIREN RetroQ vector. The IFN-inducer shRNA is specifically designed to robustly activate RIG-I signaling without targeting or silencing any endogenous human gene; a RNAPol III terminator sequence composed of a stretch of 5 thymidine residues; an *IFNB* promoter-luciferase sequence consisting of the 147 base pair long fragment of the human *IFNB* promoter containing the enhanceosome sequence (18) followed by the TATA box and the transcriptional start site, positioned just upstream of the firefly luciferase gene coding sequence. All sequences were verified at the McGill University and Genome Quebec Innovation Centre (Montreal, Canada) or at Eurofins Genomics (Ebersberg Germany).

### Antibodies

The anti-RSV NS1/NS2 antibody was generously provided by Dr. P. Collins (19). Primary antibodies used in immunoblot were anti-Flag (#F1804, Sigma-Aldrich) anti-actin (#MAB1501, Chemicon or #Sc-1616, Santa Cruz Biotechnology), anti-STAT2 (#Sc-136079, Santa Cruz Biotechnology), anti-phosphoTyr689-STAT2 (#07-224, Upstate/now Sigma-Aldrich/Merck), anti-IRF-3 (#Sc-9082, Santa Cruz Biotechnology), and anti-phosphoSer396-IRF3 (#4947, Cell Signaling Technology). Secondary horseradish peroxidase (HRP)-conjugated antibodies used for immunoblot were goat anti-mouse (31430, Thermo Fisher Scientific or 074-1806, Kirkegaard & Perry Laboratories) and goat anti-rabbit (31460, Thermo Fisher Scientific or 074-1506 Kirkegaard & Perry Laboratories).

### Cell Culture

A549 human alveolar epithelial cells (ATCC) were cultured in Ham’s F-12 nutrient mixture supplemented with 10% FetalClone III serum (FCl-III, HyClone) and 1% L-glutamine (Gibco). Unless otherwise indicated, cells were maintained without antibiotics and regularly tested for mycoplasma contamination using the MycoAlert Mycoplasma Detection Kit (Lonza). Phoenix-AMPHO packaging cells (ATCC) were cultured in Dulbecco’s Modified Eagle’s medium (DMEM) supplemented with 10% fetal bovine serum (FBS), 1% L-glutamine and 1% penicillin/streptomycin (all from Euroclone s.p.A).

### Generation and validation of A549 Cells Stably Expressing RSV Flag-NS2

A549 cell transfection was performed using the TransIT-LT1 reagent (Mirus Bio) according to the manufacturer’s protocol and as previously described (20). Stable expression of Flag-tagged NS2 (A549-Flag-NS2) was achieved by transfecting A549 cells with hNS2-3xFlag-pCMV-3Tag1a. Control cells (A549-Ctrl) were generated by transfection of the empty vector. Forty eight hours post-transfection, cells were subjected to selection with 800 µg/mL geneticin (G418, Gibco). After 10-14 days of selection, polyclonal populations were diluted to 40 cells/mL, and individual clones were isolated using cloning rings. NS2 expression in selected clones was confirmed by immunoblotting using anti-Flag antibodies. Where indicated, A549-Ctrl and A549-Flag-NS2 cells were stimulated with recombinant human IFNβ (PBL assay Science) at a concentration of 2,000 International Units (IU)/mL for the indicated times. Alternatively, subconfluent monolayers of cells were infected with Sendai virus (SeV) Cantell strain (Charles River Laboratories) at 40 hemagglutinin units (HAU)/10^6^ cells in the minimum volume of serum free medium. At 2 h post-infection, the medium was supplemented with F-12 medium and a final concentration of 10% FCl-III. The infection was pursued for the indicated time.

### Generation of A549 reporter cells stably expressing RSV NS2 and an IFN-inducer *IFNB* promoter luciferase reporter

Briefly, Phoenix-AMPHO packaging cells at approximately 80% confluence were transiently transfected after medium replacement with the pSIREN RetroQ IFN-inducer *IFNB* promoter luciferase retroviral vector using the calcium phosphate transfection Kit (Thermo Fisher Scientific) in the presence of 25 μM chloroquine (Sigma-Aldrich). The medium was changed at 16 and 24h post-transfection. After an additional culture of 24h, the supernatant containing the retroviral particles was collected and centrifuged at 377g to remove cell debris. The clarified supernatant was used to transduce A549-Ctrl and A549-Flag-NS2 cells in the presence of 8 μg/mL polybrene (Sigma-Aldrich). After 24h, the transduction medium was removed and replaced by Ham’s F-12 medium supplemented with 10% FBS, L-glutamine, and antibiotics (all from Euroclone S.p.A.). Selection of cells was performed with 1.5 μg/mL puromycin. Puromycin-resistant populations of A549-Ctrl-IFNind/*IFNB*prom-luc and A549-Flag-NS2-IFNind/*IFNB*prom-luc cells were used for downstream experiments.

### siRNA-mediated silencing of NS1 and NS2

A549 cells at 30-40% confluency in 60mm plates in antibiotic-free medium were transfected using Oligofectamine (Invitrogen) with 400 pmol small interfering RNA (siRNA) targeting RSV A2 NS1 (antisens oligonucleotide 5’-3’: UUUCACAAUUGUCAUCUAGUU) or NS2 (antisens oligonucleotide 5’-3’: “GUGACAACGGUCUCAUGUCdTdT”), as well as a non-targeting control siRNA (siCtrl) (antisens oligonucleotide 5’-3’: UGUGAUCAAGGACGCUAUGUU”) obtained from Dharmacon as previously described (21). Cells were incubated for 48 h to allow efficient siRNA-mediated silencing and were subsequently subjected to RSV infection. Briefly, cells were infected with sucrose purified RSV A2, prepared from an original stock obtained from Advanced Biotechnologies Inc, as previously described (22), at a multiplicity of infection (MOI) of 3 in F-12 medium supplemented with 2% FCl-III. The infection was allowed to proceed for the indicated time. Silencing efficiency was assessed by immunoblot analysis using the anti-RSV NS1/NS2 antibodies.

### Immunoblot analysis

For detection of NS1/NS2, whole cell extracts (WCE) were prepared in 50 mM HEPES pH 7.4, 150 mM NaCl, 5 mM EDTA, 10% glycerol, 1% IGEPAL supplemented with phosphatase inhibitors (5 mM NaF, 1 mM Na₃VO₄, 2 mM p-Nitrophenyl phosphate, 10 mM β-glycerophosphate pH 7.5, 1 mM DTT, 1 mM EGTA) and protease inhibitors (1 μg/mL leupeptin, 2 μg/mL aprotinin, 1 μg/mL pepstatin A, 1 μM PMSF) by incubation on ice for 20 min followed by 3 freeze-thaw cycles. After centrifugation at 13,000rpm at 4C for 20 min, protein concentration in the supernatant was determined using the protein assay (Bio-Rad). An equal amount of WCE (60 μg for endogenous and transient transfection; 200-300 µg for stable A549-Flag-NS2 cells) was resolved by SDS-PAGE using a 15 % acrylamide/bisacrylamide (37.5:1, Bio-Rad) gels and transferred overnight into a nitrocellulose membrane using a transfer buffer containing 20 % ethanol. Proteins were detected by immunoblotting, as detailed in (23), using anti-RSV NS1/NS2, anti-Flag and anti-actin primary antibodies followed by HRP-conjugated secondary antibodies. Immunoreactive bands were visualized by enhanced chemiluminescence (ECL) using the Western Lightning Chemiluminescence Reagent Plus (Perkin-Elmer Life Sciences) and detected using a LAS4000mini CCD camera apparatus (GE healthcare).

For STAT2 and IRF3 detection, WCE were prepared by lysis in 50 mM Tris pH 7.4, 150 mM NaCl, 1 mM EDTA pH 8, 10% glycerol, 1% NP-40, supplemented with protease and phosphatase inhibitor cocktails (Sigma-Aldrich), 5 mM NaF, and 0.05 mM β-glycerophosphate. WCE (50-300 µg) were resolved by SDS-PAGE on 10% (STAT2) or 8.5% (IRF-3) acrylamide/bisacrylamide (37.5:1, Bio-Rad) gels. Proteins were transferred to the nitrocellulose membrane for 75 min using a transfer buffer solution containing 20% methanol. Proteins were detected by immunoblotting using anti-STAT2, anti-phosphoTyr689-STAT2, anti-IRF3, anti-phosphoSer396-IRF3, and anti-actin primary antibodies followed by HRP-conjugated secondary antibodies. Immunoreactive bands were visualized by ECL using the Immobilon Western Chemiluminescent HRP substrate (Merk Millipore) and detected using a Biospectrum apparatus (UVP, Ultra-VioletProducts Ltd, Upland, CA).

### RT-qPCR

Total RNA was extracted using the RNAqueous-96 Isolation Kit (Ambion). Reverse transcription was performed with 1 µg of total RNA using the QuantiTect Reverse Transcription Kit (Qiagen) according to the manufacturer’s instructions. Quantitative PCR (qPCR) was carried out using the FastStart SYBR Green Kit (Roche) to measure transcript levels of SeV nucleoprotein gene (*N*), *IFNB*, and *TNF*, with primer sets previously described (24). Expression levels were normalized to the housekeeping gene *RPS9* (Ribosomal Protein S9). Reactions were performed on a Rotor-Gene 3000 Real-Time Thermal Cycler (Corbett Research). Standard curves for absolute quantification were generated from serial dilutions of reference plasmids, and mRNA copy numbers were calculated based on PCR efficiency and standard curve intercepts as previously described (24).

### Luciferase reporter assay in A549-Ctrl and A549-Flag-NS2 cells

A549-Ctrl and A549-Flag-NS2 cells seeded in 24-well plates 24 h prior transfection, were co-transfected with a firefly luciferase reporter plasmid driven by the *IFNB* promoter, *IFNB*prom-pGL3 described in (20), a renilla luciferase control plasmid (pRL-Null, Promega) and increasing amounts of hNS2-pcDNA5 plasmid for a total of µg DNA. Transfections were performed using the TransIT-LT1 reagent (Mirus Bio) according to the manufacturer’s instructions. At 16 h post-transfection, cells were either left untreated (mock) or infected with SeV by directly adding 10 µL of viral stock (4000 HAU/mL) to the culture medium without washing or medium replacement. Infection was carried out for 8 h prior to luciferase measurement using the Dual-Luciferase Reporter Assay System (Promega) according to the manufacturer’s instructions. Luminescence was measured using a Sunergy HT plate reader (BioTek). Firefly luciferase activity was normalized to Renilla luciferase activity for each sample and expressed as relative activities.

### Luciferase reporter assay for NS2-mediated interference in A549-IFN-ind/*IFNB*prom-luc reporter cells

A549-Ctrl-IFNind/*IFNB*prom-luc or A549-Flag-NS2-IFNind/*IFNB*prom-luc cells were lysed in 100 μl of 1X passive lysis buffer (PLB) for 15 min at room temperature before centrifugation for 2 min at 10,000 × g. For each experimental condition, 20 μl of clarified lysate were analyzed by addition of 80 μl of Luciferase Assay Reagent II (Promega) followed by luminescence measurement (30″ integration time) using a Lumat LB9501 Luminometer (E & G Berthold, Bad Wildbad, Germany). Data were presented as Relative Luciferase Units (RLU) after normalization to total protein content determined using the Protein Assay (Bio-Rad).

### Statistical analyses

Statistical analyses were performed using GraphPad Prism v5.0 for Windows (**Figure 5**) or the GraphPad Prism 10 for Mac (**Figure 1 and 2**). Statistical significance is indicated as: *p* < 0.05 (\**), p < 0.01 (*\*\**), p < 0.001 (*\*\*\*), and *p* < 0.0001 (****).

**Figure 1:**
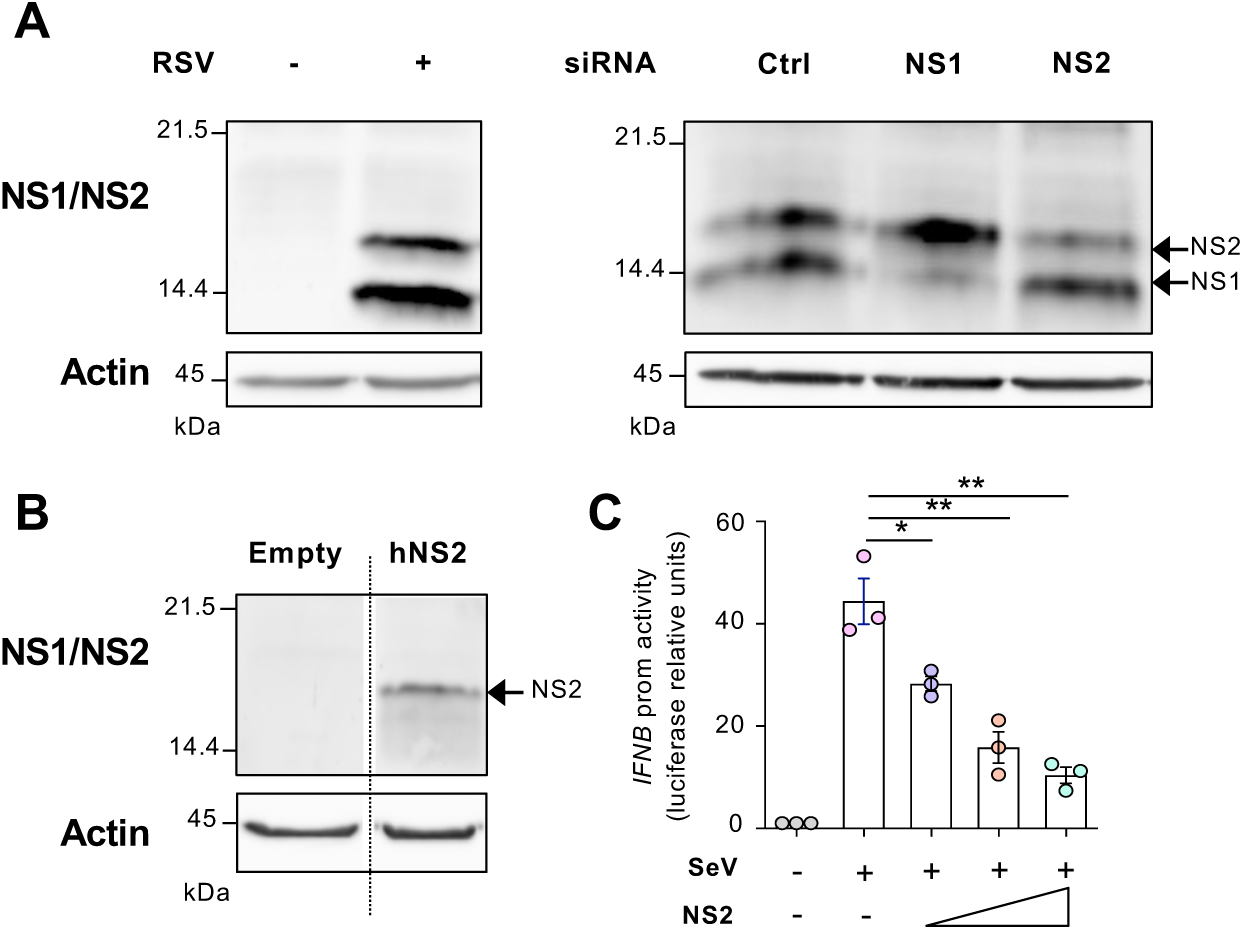
Expression of functional RSV NS2 in A549 cells. In **A)** A549 cells were infected with RSV A2 (MOI=3; 8 hrs) before analysis of WCE by immunoblotting using an anti-RSV NS1/NS2 antibody (left panel). The specificity of the anti-RSV NS1/NS2 antibody was confirmed by transfection of siRNA-targeting NS1 or NS2 before infection with RSV A2 (right panel). Anti-actin was used as a loading control. **B)** A549 cells were transiently transfected with an empty vector or the hNS2-pcDNA5 plasmid containing a humanized version of the RSV Long NS2 coding sequence and analyzed by immunoblotting using anti-RSV NS1/NS2 antibodies. **C)** A549 cells were co-transfected with an IFNBprom-pGL3 firefly luciferase plasmid, a pRL-null renilla luciferase control plasmid (pRL-Null, Promega) and increasing amounts of hNS2-pcDNA5 plasmid before infection with Sendai virus (SeV) at 80HAU/10^6^ cells. Luciferase activities were expressed as fold activation over the corresponding NS condition after normalization with renilla luciferase activities. Data (mean ±SEM; n=3) were analyzed using a two-tailed paired Student’s t-test.

**Figure 2:**
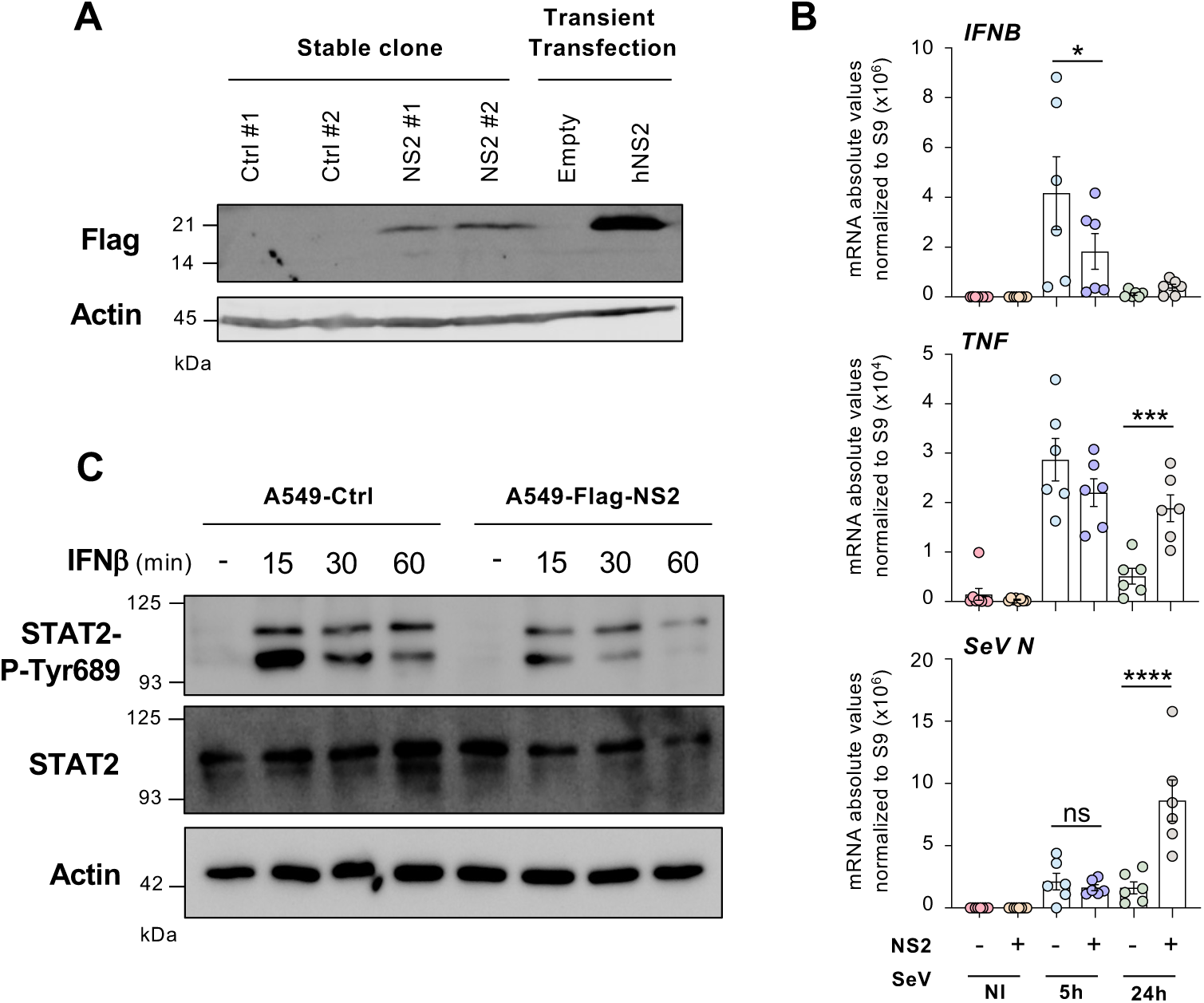
Establishment of A549 cells stably expressing functional RSV Flag-NS2. **A)** A549 cells transfected with either 3xFlag-pCMV-3Tag1a control or hNS2-3xFlag-pCMV-3Tag1a plasmids, followed by antibiotic selection to establish stable population and isolate control monoclonal cell lines (A549-ctrl) and NS2-expressing monoclonal cell lines (A549-Flag-NS2). The expression of Flag-tagged NS2 in the resulting monoclonal cells was confirmed by immunoblotting using anti-Flag antibody. Anti-actin was used as a loading control. Transiently transfected cells were used as positive control of Flag-NS2 expression. **B)** Pools of four A549-Ctrl or of A549-Flag-NS2 monoclonal cells were infected with Sendai virus (SeV) for the indicated times. IFNβ, TNF and SeV N transcript levels were quantified by RT-qPCR. **C)** A549-Ctrl and A549-Flag-NS2 cells were stimulated with recombinant human IFNβ for the indicated times. WCE were analyzed by immunoblotting using antibodies against total STAT2, phosphorylated STAT2 (Tyr689). Anti-actin was used as a loading control.

## Results

### Generation of A549 cells stably expressing RSV Flag-NS2

To establish a cellular system suitable for quantifying RSV NS2-mediated inhibition of the IFNβ response, we first generated A549 cells stably expressing NS2 from the RSV Long strain. A549 cells are a well-characterized human alveolar epithelial cell line widely used to model RSV infection, and their innate immune response reproduces key features seen in primary human airway epithelial cells (24–26). We used a previously described humanized version of the RSV Long NS2 coding sequence (hNS2), in which the AT-rich nucleotide content of the native viral gene was modified by synonymous codon substitutions to optimize expression in mammalian cells (8). Although the encoded protein sequence remains unchanged, this codon optimization significantly improves translational efficiency and has been shown to support robust NS2 protein expression in HEK293 cells (8). Thus, we first confirmed efficient NS2 expression from the hNS2-pcDNA5 plasmid by transient transfection in A549 cells and detection with anti-RSV NS1/NS2 antibodies, which we validated in parallel by siRNA knockdown of NS1 and NS2 in RSV-infected A549 cells (**Figure 1A-C**). We next demonstrated that transient NS2 expression in A549 inhibits Sendai virus (SeV)-induced *IFNB* promoter activation in a dose-dependent manner, as measured using an *IFNB* promoter luciferase reporter assay (**Figure 1D**).

To facilitate detection and support downstream biochemical and functional analyses, the hNS2 coding sequence was subcloned into the 3xFlag-pCMV-3Tag1a expression vector to generate an N-terminally Flag-tagged NS2 fusion protein. A549 cells were transfected with the resulting hNS2-3xFlag-pCMV-3Tag1a constructs, followed by antibiotic selection with G418 to establish a stable population and isolate monoclonal NS2-expressing cell lines (A549-Flag-NS2). In parallel, control monoclonal cell lines (A549-ctrl) were generated using the same protocol following transfection with the empty 3xFlag-pCMV-3Tag1a vector. Expression of Flag-tagged NS2 in the resulting monoclonal A549 lines was confirmed by immunoblotting (**Figure 2A**).

### Functionality of A549-Flag-NS2 cells

To determine whether stably expressed NS2 was functionally active and recapitulated its known antagonistic effects on the IFN responses, we assessed its ability to suppress both *IFNB* induction and downstream IFNβ signaling. A549-ctrl and A549-Flag-NS2 cells (pool of 4 clones) were infected with SeV for 5 h and 24 h, and IFNβ mRNA levels were quantified by RT-qPCR. As shown in **Figure 2B**, SeV-induced IFNβ expression was significantly reduced in NS2-expressing cells compared to control cells. This suppression was functionally relevant, as it was accompanied by a marked increase in SeV replication in NS2-expressing cells, consistent with impaired IFNβ-mediated antiviral defense. Interestingly, in contrast to its repressive effect on *IFNB* transactivation, NS2 has been reported to enhance NF-κB-dependent transcription of proinflammatory cytokine genes, such as *TNF* (19). In line with this, TNF mRNA levels were significantly elevated in A549-Flag-NS2 cells relative to control cells (**Figure 2B**). To further evaluate whether NS2 affects the cellular response to exogenous type I IFN, A549-ctrl and A549-Flag-NS2 cells were stimulated with recombinant human IFNβ for increasing duration. Total and phosphorylated (Tyr689) STAT2 levels were assessed by immunoblotting. As shown in **Figure 2C**, both total and phosphorylated STAT2 declined more rapidly in NS2-expressing cells, with a marked reduction at 60 min, supporting accelerated turnover of STAT2. Taken together, these results indicate that stably expressed NS2 retains its multifaceted immunomodulatory activity, suppressing IFNβ production, weakening antiviral responses, enhancing proinflammatory gene expression, and attenuating responsiveness to type I IFN.

### Engineering of A549-ctrl and A549-Flag-NS2 cells with an IFN-Inducer/IFNB promoter reporter System

Next, we sought to further engineer the cells to develop a virus-free, cell-based system that recapitulates NS2-mediated suppression of *IFNB* promoter activity in a controlled and reproducible manner. For this purpose, we further engineered the A549-ctrl and A549-Flag NS2 cells using a retroviral vector, pSIREN RetroQ IFN-inducer/*IFNB* promoter luciferase vector (**Figure 3**), built to enable the stable co-expression of a non-targeting (nt) short hairpin RNA (shRNA) under the control of the human U6 promoter (RNA Pol III-dependent) and a luciferase reporter gene driven by the *IFNB* promoter. The nt shRNA sequence (named IFN inducer) used in this construct does not silence any endogenous gene but was previously shown to mimic the formation of double stranded RNA species generated during active viral replication, acting as a potent activator of the RIG-I receptor and downstream *IFNB* promoter transactivation (17). Constitutive expression of the nt shRNA is thus expected to stimulate RIG-I and drive activation of the *IFNB* promoter, leading to luciferase expression. This aims to provide a direct, quantitative readout of *IFNB* promoter activity in the absence of infection (**Figure 4**).

**Figure 3:**
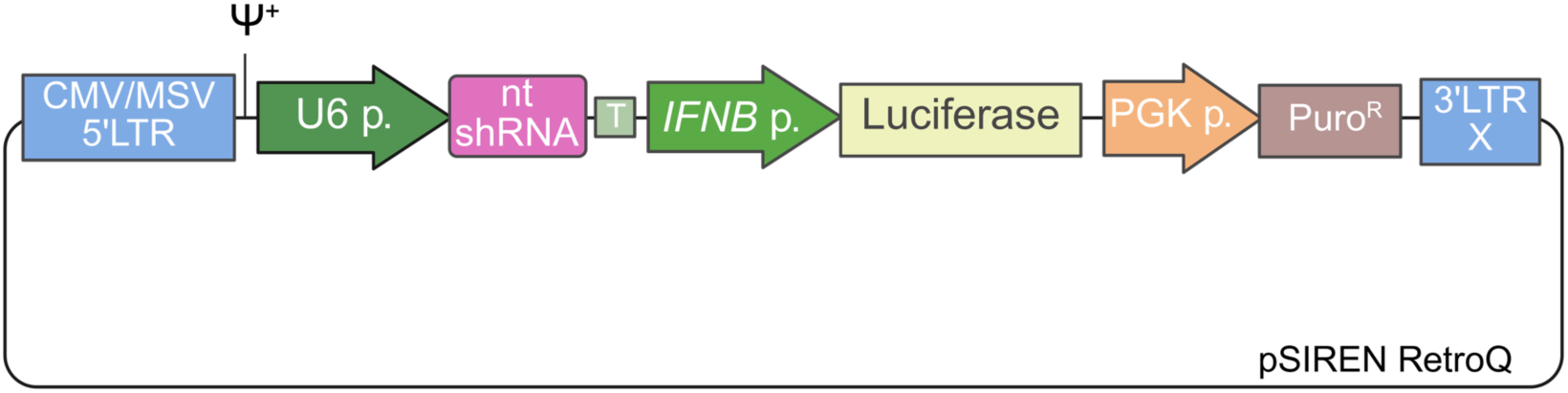
Schematic representation of the pSIREN RetroQ IFN-inducer *IFNB* promoter luciferase retroviral vector. CMV/MSV 5’LTR: hybrid 5’ Long Terminal Repeats (LTR) consisting of the cytomegalovirus (CMV) type I enhancer and the mouse sarcoma virus (MSV) promoter; Ψ+: extended packaging signal; U6 p.: RNA Pol III-dependent human U6 promoter; nt shRNA: sequence producing a non-targeting (nt) shRNA able to trigger RIG-I-dependent type I IFN production; T: RNA Pol III terminator sequence; *IFNB* p.: *IFNB* promoter sequence corresponding to the enhanceosome; Luciferase: luciferase reporter gene; PGK p.: mouse phosphoglycerate kinase 1 promoter; PuroR: Puromycin resistance; 3’LTR X: 3’ Moloney Murin Leukemia Virus Long Terminal Repeats (LTR) containing a deletion in U3 region (X) resulting in transcriptional silencing of the resulting integrated 5’ LTR.

**Figure 4:**
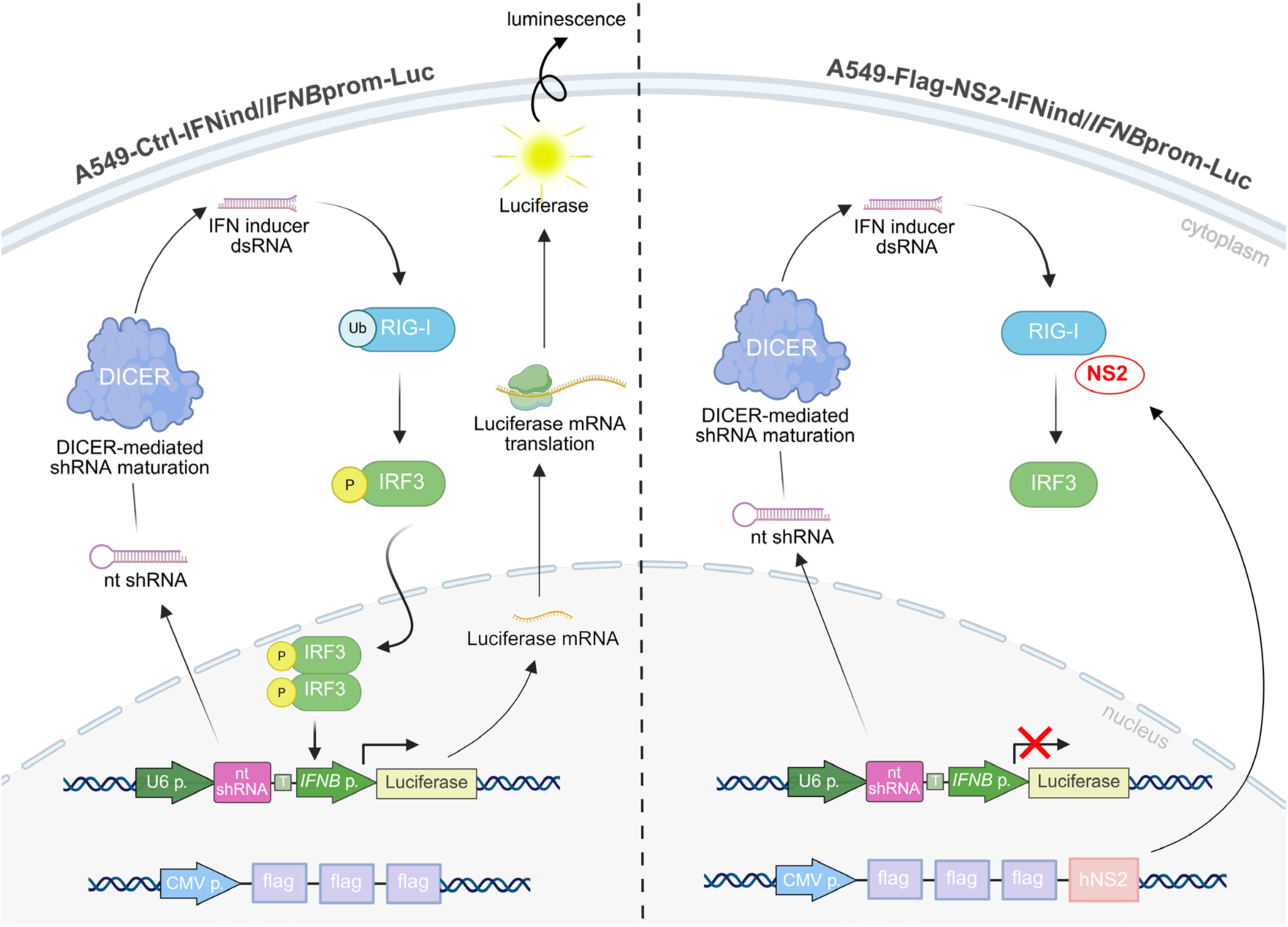
Schematic representation of the reporter cell system enabling quantitative, virus-free analysis of RSV NS2-Mediated Inhibition of *IFNB* promoter activation. A549-Ctrl-IFNind/*IFNB*prom-Luc (left) and A549-Flag-NS2-IFNind/IFNBprom-Luc (right) reporter cells were generated from stable A549-Ctrl and A549-Flag-NS2 cells, respectively, by retroviral integration of the pSIREN RetroQ IFN-inducer/*IFNB* promoter–luciferase cassette described in Figure 3. The cassette drives constitutive nuclear expression of an IFN-inducer non-targeting (nt) short hairpin RNA (shRNA) from the RNA polymerase III–dependent U6 promoter. After export to the cytoplasm, the nt shRNA is processed by the DICER machinery to generate double-stranded RNA (dsRNA) species that activate the RIG-I receptor. Activation of RIG-I, including by ubiquitination (Ub), triggers downstream signaling ultimately leading to phosphorylation and dimerization of the IRF-3 transcription factor responsible for *IFNB* promoter transactivation and luciferase induction. In A549-Flag-NS2-IFNind/*IFNB*prom-Luc cells, the expression of RSV NS2 antagonises RIG-I activation, notably through interference with its ubiquitination, thereby preventing downstream signaling and consequently reducing *IFNB* promoter–driven luciferase induction.

A549-Ctrl and A549-Flag-NS2 cells were transduced with retroviruses obtained from the pSIREN RetroQ IFN-inducer *IFNB* promoter luciferase vector and selected for stable expression of the reporter cassette. The engineered reporter cells, A549-Ctrl-IFNind/*IFNB*prom-Luc and A549-Flag-NS2-IFNind/*IFNB*prom-Luc, respectively, were first validated by confirming they conserved the NS2 expression by immunoblotting (**Figure 5A**). To confirm the functionality of the “IFN-inducer” nt shRNA construct, we next evaluated activation of IRF-3, a key transcription factor downstream of RIG-I responsible for driving *IFNB* promoter activity. IRF-3 activation was assessed by immunoblotting for phosphorylation at Ser396, a well-established marker of its activation. As shown in **Figure 5B**, strong IRF-3 phosphorylation was detected in A549-Ctrl-IFNind/*IFNB*prom-Luc reporter cells, whereas parental A549-Ctrl cells lacking the retroviral construct showed minimal signal. These results confirm that the nt shRNA efficiently activates the RIG-I signaling pathway in this system.

**Figure 5:**
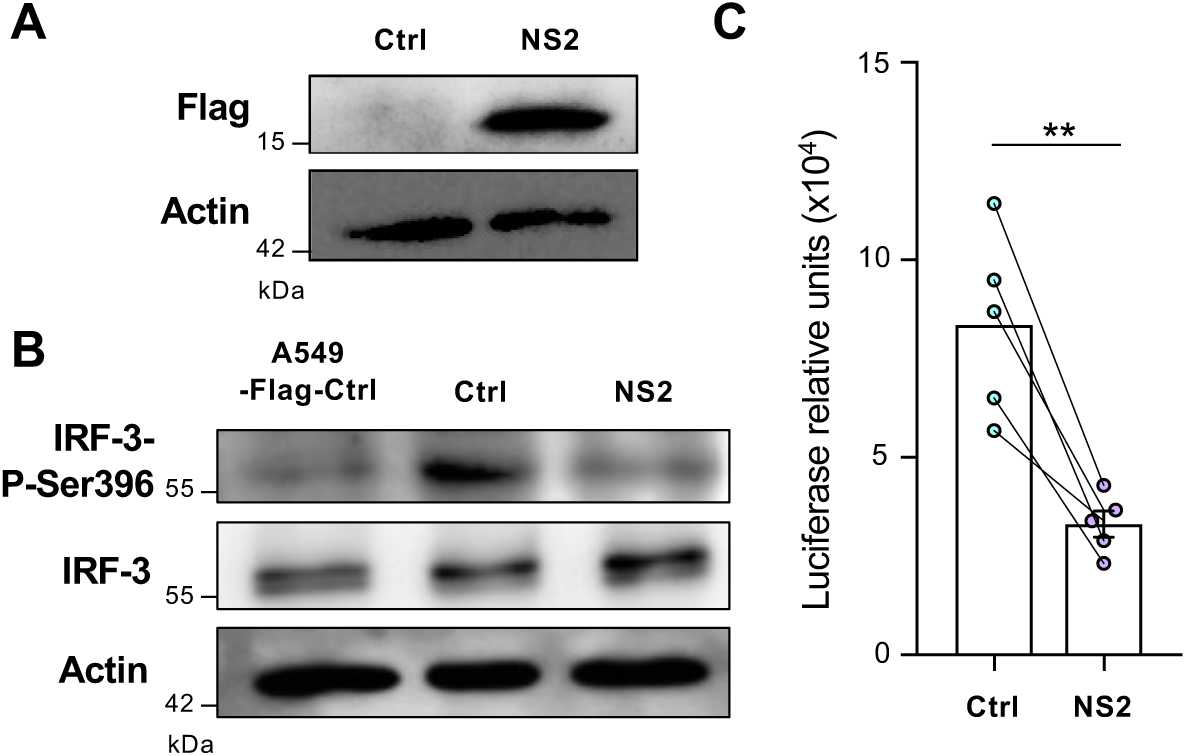
Validation of the A549-Ctrl-IFNind/*IFNB*prom-Luc and A549-Flag-NS2-IFNind/IFNBprom-Luc reporter cells. **A-B)** WCE from A549-Ctrl-IFNind/*IFNB*prom-Luc and A549-Flag-NS2-IFNind/*IFNB*prom-Luc cells were analyzed by immunoblot using anti-Flag (**A**) anti-IRF-3 and anti-IRF-3-P-Ser396 antibodies (B). Actin was used as a loading control. In **B**), A549-Ctrl cells were used as control. **C)** Luciferase activities were quantified from A549-Ctrl-IFNind/*IFNB*prom-Luc and A549-Flag-NS2-IFNind/*IFNB*prom-Luc cells and normalized over total protein content. Data (n=5 independent experiments) were analyzed using a two-tailed paired Student’s t-test.

Importantly, comparison of A549-Flag-NS2-IFNind/*IFNB*prom-Luc reporter cells with the corresponding control reporter cells revealed a marked reduction in IRF-3 phosphorylation, consistent with NS2-mediated interference in RIG-I–dependent signaling. To validate the functionality of the luciferase-based reporter system, *IFNB*prom-driven luciferase activities were compared in A549-Ctrl-IFNind/*IFNB*prom-Luc reporter cells *vs* A549-Flag-NS2-IFNind/*IFNB*prom-Luc vs reporter cells. Across five paired experiments, NS2 inhibited *IFNB*prom-driven luciferase activity by 58.9% ± 5% (SEM; vs control), a reproducible effect supported by paired analysis (paired t test p = 0.004; mean paired Δ = −50,479; 95% CI −74,643 to −26,315). This result confirms that NS2 effectively suppresses *IFNB* promoter activation in this context (**Figure 5C**). Altogether, these results demonstrate that the engineered reporter cell lines provide a reliable and sensitive readout of *IFNB* promoter activity and can be used to quantitatively assess NS2-mediated inhibition of RIG-I–dependent antiviral signaling in a virus-free setting (**Figure 4**).

## Discussion

Studying type I IFN antagonism by viral proteins such as RSV NS2 is inherently challenging. Like many viral immune antagonists, NS2 typically attenuates rather than completely abrogates signaling, yielding partial phenotypes that are highly sensitive to expression level, timing, and cellular context/compartmentalization (7, 27, 28). These features complicate the development of reliable cell-based assays to monitor interference with *IFNB* induction, because baseline pathway tone—together with experimental noise and variability—can obscure modest effects, reducing quantitative resolution and increasing the risk of ambiguous or misleading interpretations. To mitigate these limitations, we established stable A549-based reporter cell lines that enable quantitative, virus-free assessment of RSV NS2-mediated antagonism of the *IFNB* promoter activation.

A central feature of our design is an integrated reporter cassette combining an IFN-inducing nt shRNA and a luciferase reporter gene driven by the human *IFNB* promoter. Constitutive expression of the nt shRNA generates a defined double-stranded RNA species that continuously engages the RIG-I–IRF-3–IFNB axis, providing a sustained and physiologically relevant baseline of *IFNB* promoter activity, while avoiding RNAi-mediated silencing of cellular genes and the stress-associated inhibition of reporter expression sometimes observed with transient transfection (17, 29, 30). This constitutively activated, integrated system reduces sample-to-sample and experiment-to-experiment variability typically observed when *IFNB* promoter activation is driven by replicating virus or transfection of exogenous ligands. On this controlled background, the addition of stable NS2 expression further avoids key limitations of transient transfection, which encompass heterogeneous protein expression, acute cellular stress, and potential engagement of pathogen recognition receptors, such as cGAS, by transfected DNA—events that can elevate *IFNB* promoter baseline activity and suppress transgene expression—(30, 31). Overall, the stable integration of NS2 and the IFN-inducer/*IFNB*prom cassette markedly reduces well-to-well and experiment-to-experiment variability, attributes particularly important when quantifying partial antagonism. Our functional characterization confirms that stably expressed NS2 in A549 cells retains the multifaceted IFN antagonist properties described in infected cells, including suppression of IRF-3-dependent *IFNB* induction, enhancement of viral replication, STAT2 degradation, and modulation of proinflammatory gene expression. Ultimately, NS2 produced a reproducible reduction in *IFNB*prom-driven luciferase activity consistent with partial antagonism, supporting the value of a standardized stable reporter platform for resolving modest inhibitory effects.

Beyond application for dissecting signaling mechanisms in a controlled setting, the principal translational implication of our virus-free, luminescence-based readout of NS2 inhibitory activity is that it can be adapted for early-stage discovery and further scaled to high-throughput format to identify small molecules and biological modulators capable of counteracting the antagonism. Similar *IFNB*prom or ISRE-containing promoter reporter systems have successfully supported high throughput screens that identified compounds capable of restoring IFN signaling in the presence of viral antagonists or selectively suppressing virus induced *IFNB* promoter activity (17). One of the main advantages of the integrated reporter system described here is their modular nature, which allows the same experimental framework to be easily extended to other viral antagonists, signaling pathways, or cell types, for comparative studies, where pathway modulation must be evaluated under highly controlled and reproducible conditions.

We acknowledge that this system has inherent limitations. Stable overexpression of NS2 from a plasmid-based construct may not fully recapitulate the temporal dynamics, subcellular localization, or stoichiometric relationships between NS2 and other viral factors, including NS1, present during authentic RSV infection (32). Moreover, A549 cells, while valuable as a model of RSV infection of respiratory epithelium (24–26), do not capture the full complexity of the pseudostratified airway epithelium, immune cell crosstalk, or the mucosal microenvironment that influence IFN responses *in vivo*. Despite these caveats, the current A549-based reporter line provides a robust, scalable, and pragmatic system to prioritize hypotheses. By serving as a robust first-line tool for mechanistic dissection and screening, the generated data can inform and streamline subsequent studies in more physiologically relevant primary systems, such as differentiated primary airway epithelial (HAE) cultures, thereby accelerating the discovery of treatments against RSV.

## Acknowledgments

The authors thank Dr. Jacques Archambault (McGill University, Montreal, Canada) for the 3xFlag-pCMV-3Tag1a plasmid, Dr. Michael Holtzman (Washington University School of Medicine, St. Louis, Missouri, USA) for the hNS2-pcDNA5 plasmid and Dr. Peter Collins (NIH, Bethesda, Maryland, USA) for the anti-NS1/NS2 antibodies.

Figure 3 and 4 were created in BioRender. Grandvaux, N. (2026): https://BioRender.com/3dbv56g and https://BioRender.com/ fmq5bv5.

## Authors’ Contributions

LM, MS and NG contributed to the design of the study. LM, CA, MA, and DMK contributed to data collection. LM, MA, DMK, MS and NG conducted data analysis and interpretation. EGT, MS and NG wrote the original draft of the article. All authors critically revised the article and approved the final version for submission for publication.

